# CloudReg: Automatic Terabyte-Scale Cross-Modal Brain Volume Registration

**DOI:** 10.1101/2021.01.26.428355

**Authors:** Vikram Chandrashekhar, Daniel J Tward, Devin Crowley, Ailey K Crow, Matthew A Wright, Brian Y Hsueh, Felicity Gore, Timothy A Machado, Audrey Branch, Jared S Rosenblum, Karl Deisseroth, Joshua T Vogelstein

## Abstract

Quantifying terabyte-scale multi-modal human and animal imaging data requires scalable analysis tools. We developed CloudReg, an open-source, automatic, terabyte-scale, cloud-based image analysis pipeline that pre-processes and registers cross-modal volumetric datasets with artifacts via spatially-varying polynomial intensity transform. CloudReg accurately registers the following datasets to their respective atlases: *in vivo* human and *ex vivo* macaque brain magnetic resonance imaging, *ex vivo* mouse brain micro-computed tomography, and cleared murine brain light-sheet microscopy.

---

Modern imaging methods can generate intact, whole brain data from a variety of modalities including magnetic resonance imaging (MRI), computed tomography (CT), and light-sheet microscopy (LSM) of cleared tissue samples. Each of these methods provides specific information about an individual sample based on the physical principles of the technique, also producing artifacts unique to each technique. MRI can provide detailed anatomic or functional information but can be limited by intensity inhomogeneity due to magnetic field bias.^1^ CT can provide detailed anatomic information but can be limited by beam hardening.^2^ LSM, in combination with tissue clearing methods, can provide anatomic, functional, and molecular information at subcellular resolution,^3^ but can be limited by intensity inhomogeneity due to microscope optics.

Clearing methods including CLARITY (Clear Lipid-exchanged Anatomically Rigid Imaging/immunostaining-compatible Tissue Hydrogel),^3^ SHIELD (Stabilization to Harsh conditions via Intramolecular Epoxide Linkages to prevent Degradation),^4^ and iDISCO (immunolabeling-enabled three-Dimensional Imaging of Solvent-Cleared Organs)^5^ can generate terabytes of data per sample.^6^ High-resolution, multi-field-of-view (mFOV) datasets require pre-processing to remove artifacts, stitching into a complete volume, registration to a reference atlas, and visualization in order to perform quantitative analyses.^7,8^

Each of these image processing steps presents unique challenges. First, aligning and stitching every FOV acquired into a complete volume requires significant compute power and is time-intensive. Second, pre-processing imaging data requires correcting artifacts unique to each modality and sample such as intensity inhomogeneity in LSM and MRI. Third, registration methods are frequently intra-modal, have manual components, and are limited by artifacts introduced by specimen preparation and imaging.^9,10^ Finally, visualization of these terabyte-scale datasets on a local machine is compute-intensive, slow, and expensive.^11^

To address these challenges, we present CloudReg, an automatic, cross-modal, cloud-based pipeline consisting of local and global intensity correction,^12,13^ alignment and stitching,^8^ image registration with nonlinear methods,^14,15^ and interactive online visualization through Neuroglancer (https://github.com/google/neuroglancer).^16^ We specifically developed algorithms for distributed local intensity correction and cross-modal registration while leveraging existing state-of-the-art, open-source tools.^12,13^ We applied CloudReg to various datasets including *in vivo* human brain MRI,^17^ *ex vivo* macaque brain MRI,^18,19^ *ex vivo* in situ mouse brain micro-CT,^20,21^ and LSM-imaged cleared mouse and rat brains.^3,4,5^

Figure 1 shows an overview of the CloudReg pipeline. Data is uploaded to a web-accessible cloud storage provider; CloudReg is launched from a local machine (Figure 1A). The pipeline is run automatically in the cloud. CloudReg starts a cloud computing instance with sufficient RAM to perform pre-processing and registration, downloads raw data onto that server, corrects local intensity, stitches, corrects global intensity, and uploads pre-processed data to cloud storage for online visualization and analysis with Neuroglancer (Figure 1B). Next, registration is started by providing an affine initialization to roughly align an atlas to the input data and is automatically updated to include non-linear deformation via an expectation-maximization optimization process. Then, the atlas anatomic parcellations are automatically transformed to the input data at high resolution for visualization (Figure 1C). All the data is stored in the cloud, and then routed through a content delivery network and firewall to facilitate efficient and secure visualization via Neuroglancer (Figure 1D). The resulting data can be shared by sending a Universal Resource Locator (URL) and visualized from anywhere with a web browser and internet connection (example URL in Supplemental Note 1). Our deployment of Neuroglancer enables instant visualization of multi-channel, terabyte-scale datasets and provision of a shortened URL with one-click to share data views and analysis results (Supplemental Figure 1).^16^

**Figure 1.**
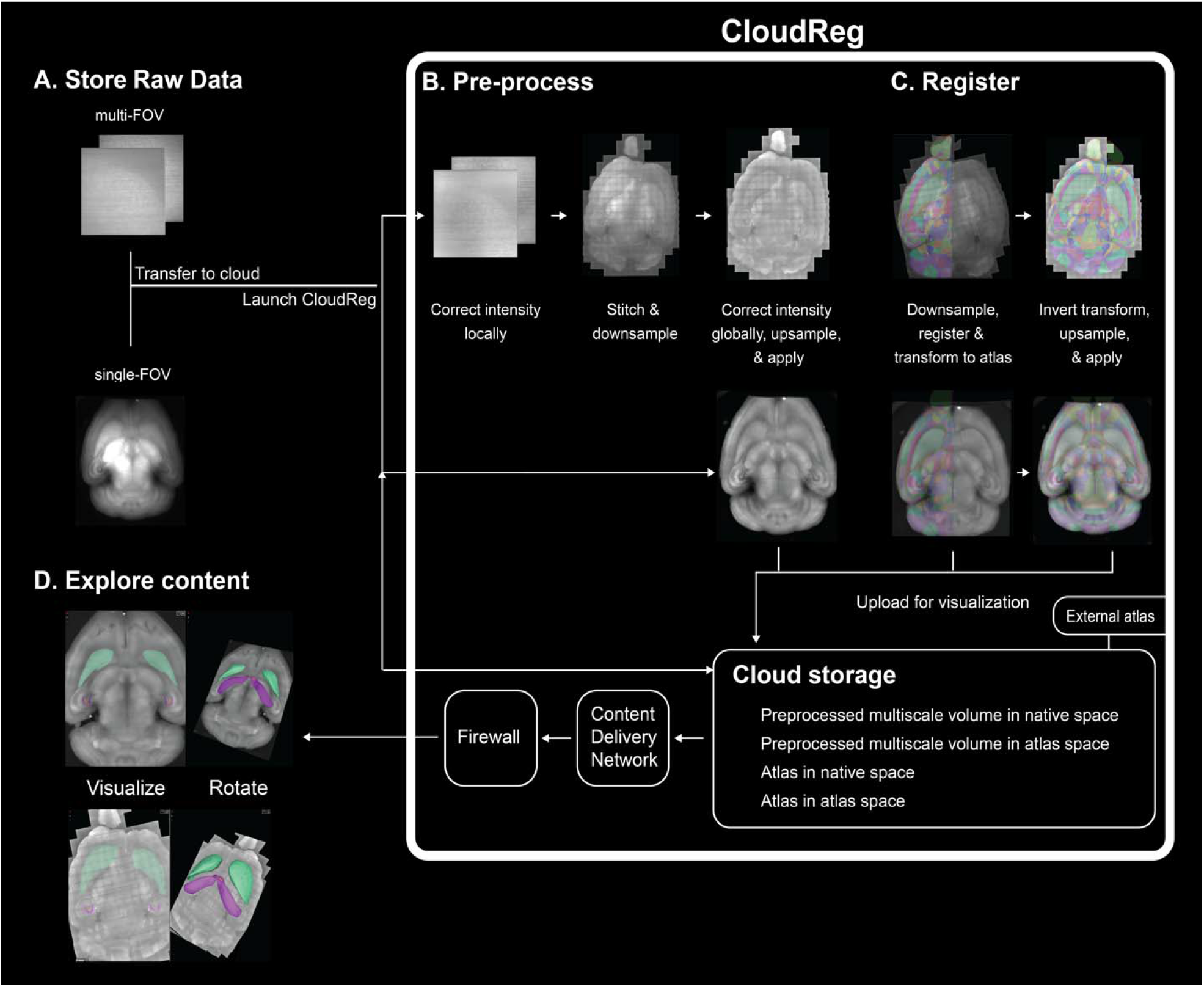
CloudReg pipeline schematic, example outputs at each step. A: Store raw data. Raw data can be either multi-field-of-view (mFOV) or single-field-of-view (sFOV). After raw data is generated, it is transferred to the cloud and CloudReg is launched. *B:* Pre-process. All of the following steps are performed on a cloud compute server that has access to the raw data. First, local intensity inhomogeneity is corrected per-FOV using an algorithm we developed. Next, mFOV data is aligned and stitched into a complete volume using Terastitcher; if the data is sFOV, this step is not performed. Then, global intensity inhomogeneity is corrected on the stitched whole brain by computing an intensity correction on the downsampled data, upsampling the result, and applying it at native resolution; this step applies to both mFOV and sFOV data. The pre-processed data is then downsampled and uploaded to cloud storage for visualization.
*C:* Register. A downsampled version of the pre-processed data is registered to the ARA CCFv3 and the computed transformations are saved. These transformations are invertible so the atlas can be transformed to the data space and the data can be transformed to the atlas space. The transformations are upsampled and applied to the ARA anatomic parcellations, and input data and are uploaded to cloud storage for visualization through Neuroglancer. To maintain privacy of imaging data, visualization is restricted to authorized users by using a content delivery network and firewall. *D:* Explore content. Left column shows ARA parcellations transformed to the s-and mFOV data based on the computed transformations from the registration. Right column shows 2D axial slice from the pre-processed input data and 3D rendering of Caudoputamen and Hippocampal regions selected from the transformed ARA visualized in Neuroglancer. ARA CCFv3, Allen Reference Atlas Common Coordinate Frame version 3.

In the next two paragraphs, we elaborate on the pre-processing and registration steps (Figure 1B and 1C, respectively). We initially developed CloudReg using high-resolution, LSM-imaged CLARITY mouse brain data^6^ and used the Allen Reference Atlas (ARA) Common Coordinate Frame Version 3 (CCFv3) as the reference atlas (Figure 2, rows 1 and 2).^22^

**Figure 2.**
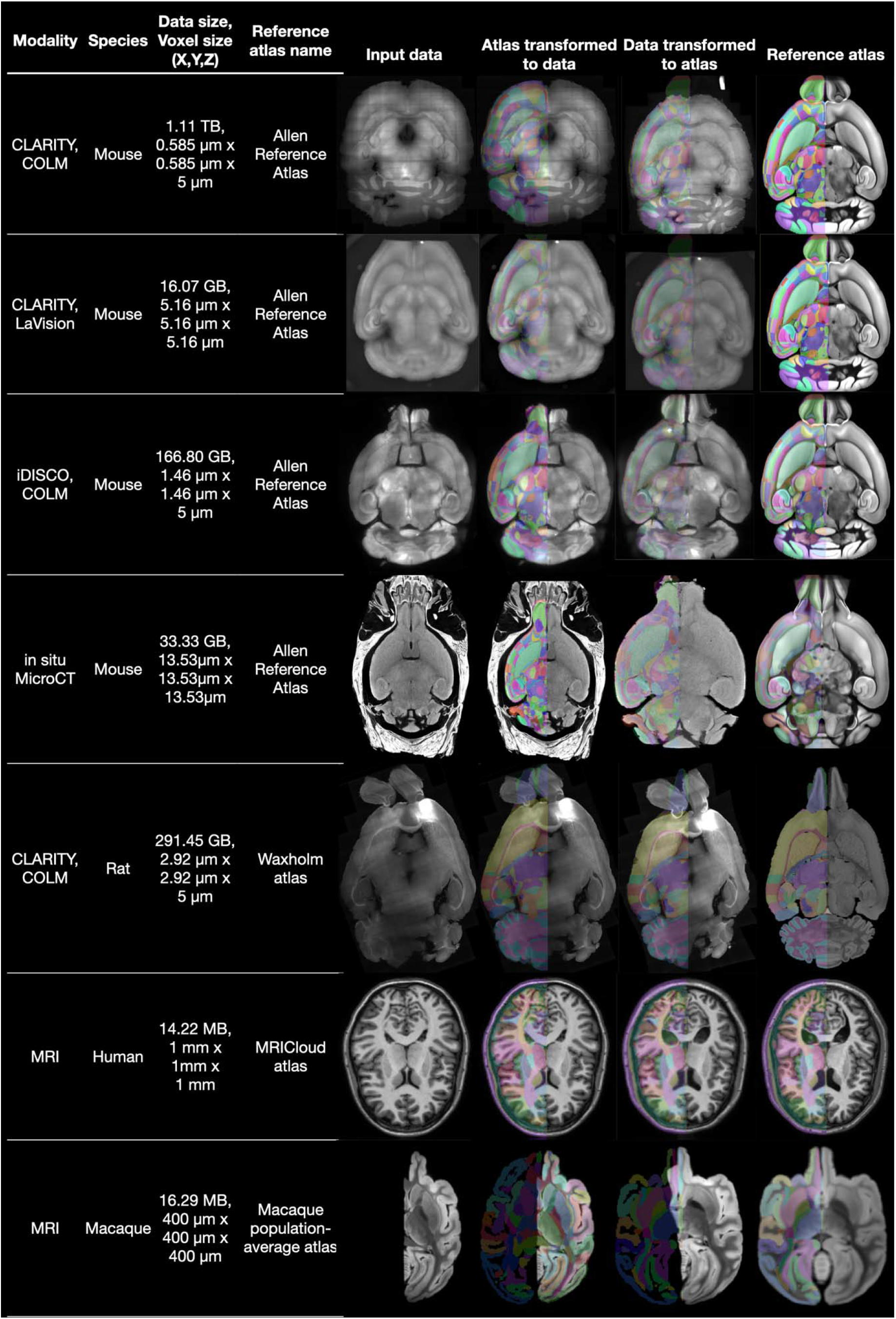
CloudReg pipeline registration outputs from multiple species imaged with various modalities. Each row demonstrates registration of either mouse, rat, macaque, or human brain imaging data to the corresponding atlas using CloudReg. The leftmost column of images shows the input data; the data from the autofluorescence channel is used for samples imaged with a light-sheet microscope (LSM). The rightmost column shows the atlas parcellations overlaid on one hemisphere of the atlas image data. The second and third columns show the respective atlas parcellations transformed to and overlaid on the original samples and vice-versa, respectively. CLARITY, Clear Lipid-exchanged Anatomically Rigid Imaging/immunostaining-compatible Tissue Hydrogel; COLM, CLARITY-Optimized Light-sheet Microscopy; GB, Gigabyte; iDISCO, immunolabeling-enabled three-dimensional imaging of solvent-cleared organs; MB, Megabyte; Micro-CT, Micro-Computed Tomography; TB, Terabyte.

Tissue clearing procedures and optics of LSM introduce sample-specific artifacts and intensity inhomogeneity in the imaged samples, which we aimed to correct with our mFOV-based pre-processing algorithm. Intensity inhomogeneity manifests as a decay of intensity away from the center of each FOV and, for mFOV samples, the center of the stitched whole brain. This makes automatic, intensity-based registration a significant challenge. To minimize this per-FOV artifact, we developed a parallelized intensity correction algorithm based on the hypothesis that the introduced intensity inhomogeneity is the same in each FOV. To efficiently compute this correction, we uniformly subsample the mFOV data in three dimensions, compute the mean across subsampled FOVs in parallel, and apply the N4 bias correction algorithm^12^ to the resulting mean FOV (Supplemental Figure 2A). Our intensity correction algorithm accounts for differences in tissue scattering from different clearing methods by estimating the intensity correction directly from the data. This pre-processed data is then automatically aligned and stitched using Terasticher, an open-source tool for stitching teravoxel microscopy datasets.^8^ To minimize intensity inhomogeneity at the whole-brain scale, we apply the N4 bias correction algorithm to the whole stitched volume directly (Supplemental Figure 2B-C).

The fully pre-processed sample, which can be acquired from a variety of modalities in a number of species, is then registered to a corresponding reference atlas. To enable registration of LSM-imaged tissue samples, we developed a spatially-varying polynomial intensity transform, expanding the scope of samples that can be automatically registered (Figure 2; Supplemental Figure 3, Supplemental Video 1). CloudReg computes affine and nonlinear transformations by building on the Expectation-Maximization Large Deformation Diffeomorphic Metric Mapping (EM-LDDMM) registration algorithm that we previously developed.^15^ Our extension of EM-LDDMM built into CloudReg enables cross-modal registration of a diversity of brain volume samples with artifacts, tears, and deformations (Supplemental Figure 4).

To assess registration accuracy, we used Target Registration Error (TRE) by computing the Euclidean distance between 19 landmarks (Supplemental Figure 5) placed by experts on the ARA and our samples, where possible. TRE for samples 1, 2, and 3 was 2.27 ± 0.86 voxels, 2.42 ± 1.31 voxels, and 2.74 ± 0.88 voxels, respectively (Supplemental Table 1). These voxels are 100 μm indicating the resolution at which the registration was performed. Measurements and summary statistics can be found in Supplemental Table 1. Examining landmark error by brain region shows regional error differences, reflecting gross regional deformations. Mean landmark error for midbrain, cortical, and cerebellar regions across all samples are 1.47, 2.32, and 2.65 voxels, respectively. The region with the greatest error, the cerebellum, was most often subject to gross deformation including rotation and translation relative to the rest of the brain. By comparison, the midbrain, a relatively fixed structure in the brain and scan, had the lowest error. However, we found that the midbrain is most subject to the two artifacts typical of tissue clearing and LSM methods: intensity inhomogeneity and hydrogel-based deformation. Our TRE demonstrates that CloudReg handles artifacts typical of hydrogel-based tissue clearing methods including intensity inhomogeneity, hyperlocal structural deformation (nonuniform, micron-scale deviation from true anatomic position), and local missing tissue exceedingly well. CloudReg achieves this by relying on a rough affine initialization. Thus, in gross regions of the brain that have been displaced relative to the rest of the intact brain, the error will increase.

Registration is a crucial first step in analyzing a single or cohort of samples but can be more informative if combined with additional downstream analysis methods including cell and axon detection. A potential extension of our current work will be to accelerate the registration component of the code by leveraging our existing C++ implementations (https://github.com/InsightSoftwareConsortium/ITKNDReg).

CloudReg can correct intensity, align and stitch, register, and visualize terabyte-scale brain volumes with artifacts and tears. CloudReg is immediately applicable to brain volumes spanning a variety of species and imaging modalities including mouse, rat, monkey, and human brain imaging.

## Methods

All methods and additional supplementary information are available in the online e-methods.

## Acknowledgments

The authors would like to thank NeuroNex and Microsoft Research for supporting this work.

## Author contributions

VC conceived the study, drafted the original manuscript, performed data analysis, built reproducible pipeline, revised the manuscript. DJT conceived the study, revised the manuscript, performed data analysis, and supervised data analysis. DC ported registration algorithm to Python, performed data analysis, and revised the manuscript. AKC, MAW, BYH, FG, TAM generated CLARITY/iDISCO LSM data and revised the manuscript. JSR helped draft the original manuscript, revised the manuscript, performed experiments, generated data, and placed landmarks for determining registration accuracy. KD conceived the study, supervised data analysis, and revised the manuscript. JTV conceived the study, revised the manuscript, and supervised data analysis.

## Competing interest statement

The authors declare no competing interests.

## Notes

https://cloudreg.neurodata.io

### Competing Interest Statement

The authors have declared no competing interest.

