## Supplement for "CloudReg: Automatic Terabyte-Scale Cross-Modal Brain Volume Registration"

**Supplemental Materials**

**Methods…2-10**

**Supplemental Note…11**

**Supplemental Figures…12-19**

**Supplemental Video…20**

**Supplemental Table…21-22**

### Cleared brain specimen preparation and image acquisition

CloudReg was developed on multiple brain imaging modalities including a variety of clearing methods, all imaged with light-sheet microscopy (LSM). Whole mouse and rat brains were optically cleared using CLARITY (Clear Lipid-exchanged Anatomically Rigid Imaging/immunostaining-compatible Tissue hYdrogel), SHIELD (Stabilization to Harsh conditions via Intramolecular Epoxide Linkages to prevent Degradation), or iDISCO (immunolabeling-enabled three-Dimensional Imaging of Solvent-Cleared Organs) as previously described.[3,4,5] Autofluorescence image volumes were acquired using either a CLARITY-Optimized Light-sheet Microscope (COLM)[6] or a LaVision UltraMicroscope II. We used the autofluorescence channel to register, and then applied the resulting transformation to any additional channels, but any/all channels could be run through CloudReg. The COLM imaged whole mouse and rat brains with voxel size 0.585x0.585x5.0 $\mu m^3$, 1.46x1.46x5.0 $\mu m^3$, or 2.9x2.9x5.0 $\mu m^3$ resulting in terabytes of data per brain for these higher resolution samples. The LaVision UltraMicroscope II was used to acquire a whole mouse brain in a single z-stack at 5.16 μm isotropic resolution.

### *Ex vivo* in situ mouse brain micro-computed tomography

The intact mouse head was imaged via micro-computed tomography (micro-CT) following terminal vascular Microfil® polymer perfusion as previously described.[20] Soft tissue including brain was visualized by immersing the sample in phosphotungstic acid (PTA) before micro-CT as previously described.[21]

### *In vivo* human brain magnetic resonance imaging

Human brain magnetic resonance imaging (MRI) data was obtained from the MRICloud atlas set as previously described.[17]

### Upload raw data to cloud storage

To run CloudReg, raw data is made web-accessible, for example, by uploading to cloud storage. We use Amazon Web Services (AWS) Simple Storage Service (S3) for our cloud storage services. The multi-field-of-view (mFOV) raw data was stored as a 2D TIFF (Tagged Image File Format) series for each column in the image volume. mFOV data was organized in the COLM acquisition format,[6] but any format can be used with minor modifications. The single-FOV (sFOV) raw data was stored as a 2D TIFF series for each slice in the image. The raw data was uploaded to S3 using the awscli Python package (https://github.com/aws/aws-cli).

### Run CloudReg

CloudReg consists of two user-facing Python scripts that we developed along with other utility functions (available here: https://cloudreg.neurodata.io). All scripts referred to herein are available in this repository. Our two user-facing scripts automatically start and stop a cloud server after running a series of computations on that server. The first script requires a cloud server with attached solid-state drives (SSD) and the second script requires a server with sufficient memory for the given data. An AWS cloud server is called an Elastic Compute Cloud (EC2) instance. The first script (`run_colm_pipeline_ec2.py`) automatically starts an r5d-type EC2 instance with attached SSD storage and performs the following steps (Figure 1A-B): 1) transfers raw data from cloud storage, 2) corrects local intensity, 3) stitches the data into a three-dimensional volume, 4) corrects global intensity, and 5) downsamples and uploads stitched and preprocessed data to cloud storage for visualization. The second script (`run_registration_ec2.py`) automatically starts an r5-type EC2 instance and performs the following steps (Figure 1C-D): 1) downloads downsampled, preprocessed data and the reference atlas data from cloud storage, 2) registers input data to the provided atlas, 3) transforms atlas anatomic parcellations to input data space and input data to atlas space, and 4) uploads transformed data to cloud storage for visualization.

For example, for mouse brain data we use the Allen Reference Atlas (ARA) Common Coordinate Frame Version 3 (CCFv3), but any reference atlas or sample can be used.

**Transfer raw data from cloud storage**

After the cloud server is started, available SSDs must be formatted and mounted onto the server for use. We developed a bash script (`mount_combined_ssds.sh`) to do this automatically for any EC2 instance with SSD storage available. The raw data is then downloaded from cloud storage onto these SSDs. Data download is parallelized across all available cores to speed up the process via a Python script (`download_raw_data.py`).

**Correct local intensity**

Our local intensity correction can be applied to any mFOV samples with intensity inhomogeneity, though we developed our local intensity correction on LSM-imaged CLARITY data. For mFOV data, there are two intensity correction steps: 1) local correction per FOV and 2) global correction on the stitched image volume. A single intensity correction step is performed on on the whole image volume of sFOV data. To correct the per FOV intensity inhomogeneity, we developed an algorithm to estimate a multiplicative intensity correction directly from the data. Our algorithm begins with sub-sampling the mFOV raw data such that FOVs are uniformly sampled in all three dimensions. The amount of subsampling is a configurable parameter; increased subsampling can speed up the rest of the computation but may provide a less accurate estimate of the intensity correction. We use a subsampling factor of two. Next, the mean is computed across these uniformly subsampled FOVs such that the resulting mean is a 2D image. The multiplicative intensity correction is then estimated by applying the N4 bias correction algorithm to the computed mean image.[10,11] This algorithm is available as a Python script (correct_raw_data.py). Example results from this local intensity correction are shown in Supplemental Figure 1.

**Stitch mFOV data into a volume**

Stitching mFOV data into a volume was performed using Terastitcher,[8] an open-source software for stitching terascale images. Terastitcher uses a maximum intensity projection, normalized cross correlation (MIP-NCC) method to align each of the individual 2D FOVs in the 3D volume. Generating accurate stitching results requires each 2D FOV to have some overlap with neighboring FOVs. The stitched image volume is saved as a 2D TIFF series on the SSDs of the r5d-type EC2 instance. To accelerate this process, we adapted Python scripts from Terastitcher to parallelize stitching across all available cores subject to memory constraints. To use these scripts with our pipeline, we had to first convert them to Python 3 (paraconverter.py and parastitcher.py). Functionality in these scripts is wrapped and available in a Python script (stitching.py).

**Upload**

Stitched data is uploaded to cloud storage, for example S3, in a highly parallelized fashion using cloud-volume (https://github.com/seung-lab/cloud-volume), an open-source Python package that can write Neuroglancer-compatible data to cloud storage. We have made contributions to cloud-volume to provide additional compression methods (https://github.com/seung-lab/cloud-volume/pull/291). The stitched data is concurrently downsampled using a package called tinybrain (https://github.com/seung-lab/tinybrain) for quicker visualization and uploaded to cloud storage. We implemented this parallelized upload procedure to leverage as many cores as are available subject to memory constraints (create_precomputed_volume.py).

**Correct global intensity**

To correct intensity inhomogeneity in the stitched image volume, we applied the N4 bias correction algorithm to a downsampled version of the whole volume in 3D.[12] The downsampled intensity correction produced is then upsampled and applied to the native resolution data, which is then uploaded back to cloud storage in Neuroglancer precomputed format on a slice-by-slice basis (`correct_stitched_data.py`). Example results from this global intensity correction are shown in Supplemental Figure 1.

**Visualize with Neuroglancer**

Once the data is uploaded to cloud storage, we use Neuroglancer to view that data in a web browser. To enable sharing data/visualization by web link, we implemented a custom JSON-state server with our Neuroglancer deployment (https://viz.neurodata.io). To enable visualization of multi-channel LSM data, we added differential channel coloring that has been merged into the Neuroglancer core code base.[16] We also set up a Content Delivery Network through AWS called CloudFront on top of our S3 bucket to enable HTTP/2 to speed up visualization on Neuroglancer. We also added Internet Protocol (IP)-address-based restriction to ensure limited access to private data using AWS Web Application Firewall (WAF).

**Register to a reference atlas**

CloudReg computes affine and nonlinear transformations using a modified version of the Expectation-Maximization Large Deformation Diffeomorphic Metric Mapping (EM-LDDMM) registration algorithm.[15] Our modified EM-LDDMM enables cross-modal registration by estimating spatially-varying polynomial transformations of the atlas intensity to match our input data intensities. Per-voxel error signals in our intensity transform allow detection of artifacts and missing tissue. These concepts are combined within an EM framework where deformation parameters and polynomial coefficients are updated iteratively.

Below is a description of each variable necessary to specify an objective function to be optimized for image registration.

| Name | Definition | To be optimized |
|---|---|---|
| $x$ | A point in 3D space describing the location of a voxel. | |
| $J(x)$ | The input image which is a real-number-valued function of $x$ | Input; Fixed parameter |
| $I(x)$ | The atlas image which is a real-valued function of $x$ | Input; Fixed parameter |
| $v(x)$ | A velocity field which is a 3D vector-valued function of $x$ | Yes |
| $\varphi(x)$ | A position field which is a 3D vector-valued function of $x$. It includes a nonlinear component found from integrating $v(x)$ and an affine component. | Yes |
| $\tilde{I}(x)$ | $(1, I, I^2, I^3)^{\mathrm{T}} \circ \varphi^{-1}(x)$ | Yes |
| $c(x)$ | A 4D vector-valued function of x representing the coefficients of a 3rd order polynomial contrast transform at each voxel in $J(x)$ | Yes |
| $w(x)$ | A real-number-valued function of $x$ taking values between 0 and 1 representing the posterior probability that a voxel in $J(x)$ corresponds to some voxel in $I$ as opposed to missing tissue or artifact | Yes |
| $\sigma_M$ | A positive real number representing the standard deviation of the noise in the image $J$ and a weighting of the matching term in our objective function. To calculate $w$ we assume background and artifact has twice and five times the standard deviation, respectively. | Fixed parameter; unitless; default is standard deviation of $J$ |
| $\sigma_R$ | A positive real number representing the weighting of the regularization of $v$ in our objective function. | Fixed parameter; unitless; default is 10,000 |
| $\sigma_C$ | A positive real number representing the weighting of the regularization of $c$ in our objective function | Fixed parameter; unitless; default is 5 |
| $a$ | A characteristic length scale for regularizing $v$ | Fixed parameter; microns; default is 500 |
| $a'$ | A characteristic length scale for regularizing $c$ | Fixed parameter; microns; default is 750 |
| $L_a/L_{a'}$ | A highpass differential operator for encouraging spatial smoothness in regularization equal to $(1 + a^2\Delta)^2$ where $\Delta$ is the Laplacian. | |
| $id$ | Identity operator | |

The objective function we minimize is

$$\underbrace{\frac{1}{2\sigma_M^2}\int |c^T(x)\hat{I}(x) - J(x)|^2 w(x)dx}_{\text{Matching accuracy}} + \underbrace{\frac{1}{2\sigma_R^2}\int_0^1\int |L_a v_t(x)|^2 dxdt}_{\text{Deformation regularization}} + \underbrace{\frac{1}{2\sigma_C^2}\int |L_{a'}c(x)|^2 dx}_{\text{contrast regularization}}$$

Note that $c^T\hat{I}$ is the atlas deformed and intensity-transformed to the input data. The highpass operator used in regularization of $v$ and $c$ were initially described in Beg et al.[14] $w$ is a set of weights estimated using the EM algorithm, designed to downweight voxels that contain missing tissue or large artifacts. For a given $w$, the cost is optimized over $v$, $c$, and affine parameters using gradient descent. This is the maximization step of the EM algorithm. For the expectation step, $w$ is updated using Gaussian mixture modeling (GMM) which depends on the value $\sigma_M$. The procedure is described in more detail in Tward et al.[15] Our contribution in this work is twofold: (1) we introduce including $c$ as a function of space rather than a constant, and (2) we add a contrast regularization term in the objective function.

CloudReg can be run on any two image volumes that have correspondence. We optimized aspects of the pipeline for mouse and rat, whole-brain, LSM-imaged CLARITY data. Specifically, registration is performed on a downsampled version of the input data, at 100 μm. The resulting transformations are upsampled and applied to the atlas to transform it to the input data coordinate space and the input data to the atlas space. The resulting transformations generated by EM-LDDMM are smooth and can be upsampled with linear interpolation,[15] enabling visualization of the registered atlas at the native resolution of our observed image. Transformations are stored on cloud storage with transformed images stored in Neuroglancer precomputed format for visualization. Our modified EM-LDDMM is available in CloudReg as a MATLAB script (`map_multiscale_nonuniform_v02_mouse_gauss_newton.m`) and exists in a fork of scikit-image, an open-source python image analysis toolkit (https://github.com/scikit-image/scikit-image/pull/4390). The python script remains four-fold slower than the MATLAB script largely due to the relative computational efficiency of 3D interpolation.

**Spatially varying polynomial intensity transform**

Since the objective function we optimize (see equation above) is quadratic in $c$, we can solve for $c$ by solving a linear system of equations, given by

$$\frac{1}{\sigma_C^2} L_{a'} L_{a'} c + \frac{1}{\sigma_M^2} \hat{I}\hat{I}^T w c - \frac{1}{\sigma_M^2} \hat{I} w J \; = \; 0$$

In the above expression we use the fact that $L$ is self-adjoint. Since the first two terms on the left correspond to positive semidefinite operations, we can rewrite the problem as:

$$\mathrm{B} = \frac{\hat{\mathrm{I}}^{\mathrm{T}}\sqrt{\mathrm{w}}}{\sigma_M} \qquad \mathrm{A} = \frac{L_{a'}}{\sigma_C} \qquad \mathrm{b} = \frac{\hat{I} w J}{\sigma_M^2}$$

$$AAc + B^T Bc - b = 0$$

To improve conditioning, and using the fact that $A$ is easily inverted in the Fourier domain because it is diagonalized by the Fourier transform, we use an elimination approach, letting $y = Ac$, and solving the system:

$$(id + A^{-1} B^T B A^{-1}) y - A^{-1} b = 0$$

by subtracting a constant multiplied by the residual (the left-hand side of the above equation) at each step (*i.e.,* gradient descent). The code to solve this system and compute the coefficients $c$ is available in a MATLAB script (`estimate_coeffs_3d.m`).

**Determine registration accuracy**

To determine registration accuracy, an expert placed 19 landmarks shown in Supplemental Figure 3 on both the ARA and three input LSM-imaged cleared tissue samples. In areas of significant deformation or damaged tissue, landmarks were not placed. The transformations computed in the registration are applied to the landmarks placed on the LSM-imaged cleared tissue data to transform them to the landmarks placed on ARA data (Supplemental Figure 5). The Euclidean distance between the LSM-imaged cleared tissue transformed points and corresponding ARA points is computed and reported in microns in Supplemental Table 1. We developed a Python script that computes registration accuracy given two Neuroglancer links with landmarks and computed transformations (`registration_accuracy.py`).

### Statistical analysis

We computed the mean and standard deviation of landmark error across all landmarks for each of our three LSM-imaged cleared tissue samples and computed means across landmarks by region including cortex, midbrain, and cerebellum.

### Utility Functions

We have also implemented a Python script to downsample and upload a 3D image stack to the cloud for visualization with Neuroglancer (`ingest_image_stack.py`).

**Supplemental Note 1**

https://ara.viz.neurodata.io/?json_url=https://json.neurodata.io/v1?NGStateID=eWgRPzTxJsNmew

**Supplemental Note 1: Allen Reference Atlas visualized with Neuroglancer.** The Neuroglancer URL (Universal Resource Locator) above shows the Allen Reference Atlas (ARA) Common Coordinate Frame version 3 (CCFv3) serial two-photon tomography raw data with anatomic parcellations overlaid and rendered in 3D.

### Supplemental Figures

**Supplemental Figure 1**

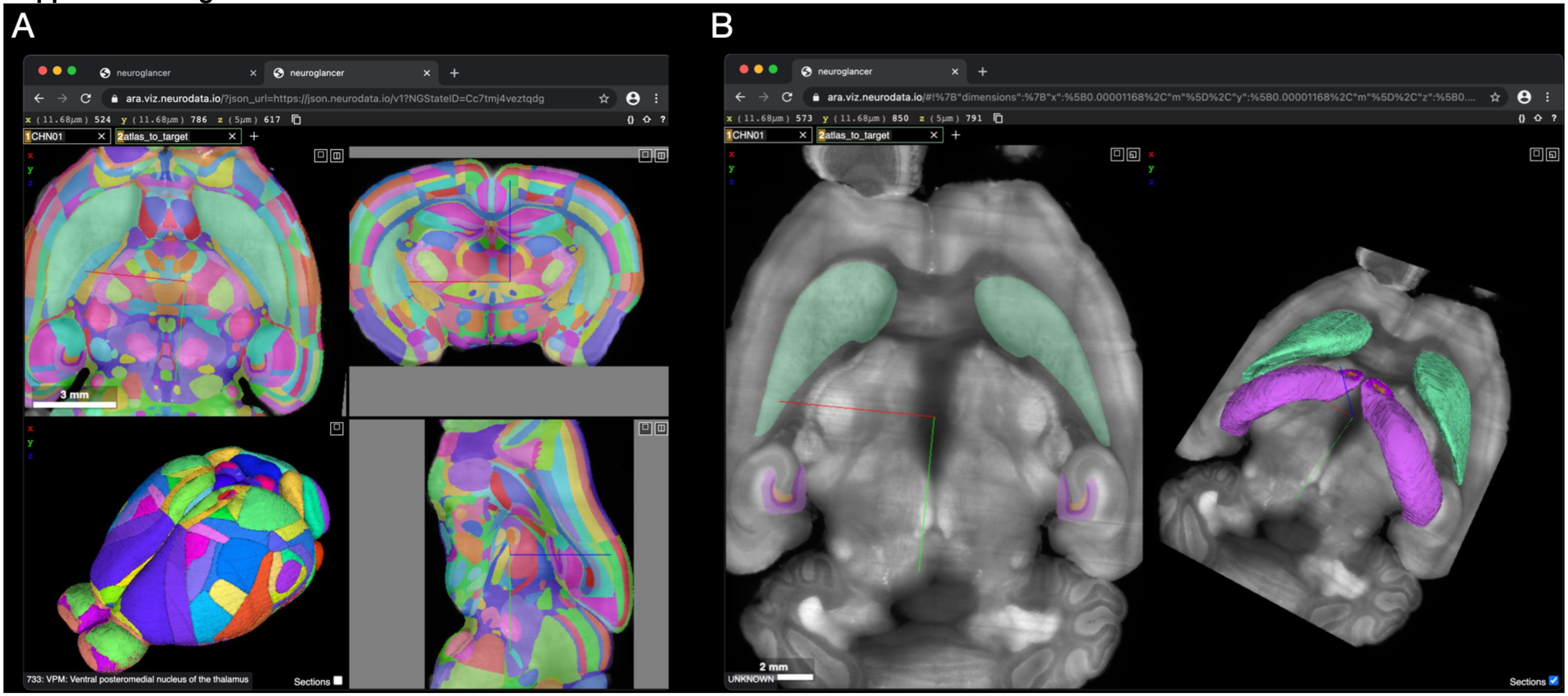


**Supplemental Figure 1: Interactive web-based visualization with Neuroglancer.** *A:* CLARITY mouse brain with ARA parcellations overlaid. LSM-imaged CLARITY-cleared mouse brain with the ARA CCFv3 anatomic parcellations registered and transformed to the input sample is shown. A 3D rendering of the resulting ARA brain nuclei segmentation based on our registration is shown in the bottom left quadrant. This data is being visualized in a web browser and is served from cloud storage. *B:* CLARITY mouse brain with selected ARA regions. The left side shows the sample from A with only Caudoputamen and Hippocampal regions selected from the ARA. The right side shows those regions rendered in 3D and overlaid on the raw data. CLARITY, Clear Lipid-exchanged Anatomically Rigid Imaging/immunostaining-compatible Tissue Hydrogel; ARA CCFv3, Allen Reference Atlas Common Coordinate Frame version 3; LSM, Light-Sheet Microscopy.

**Supplemental Figure 2**

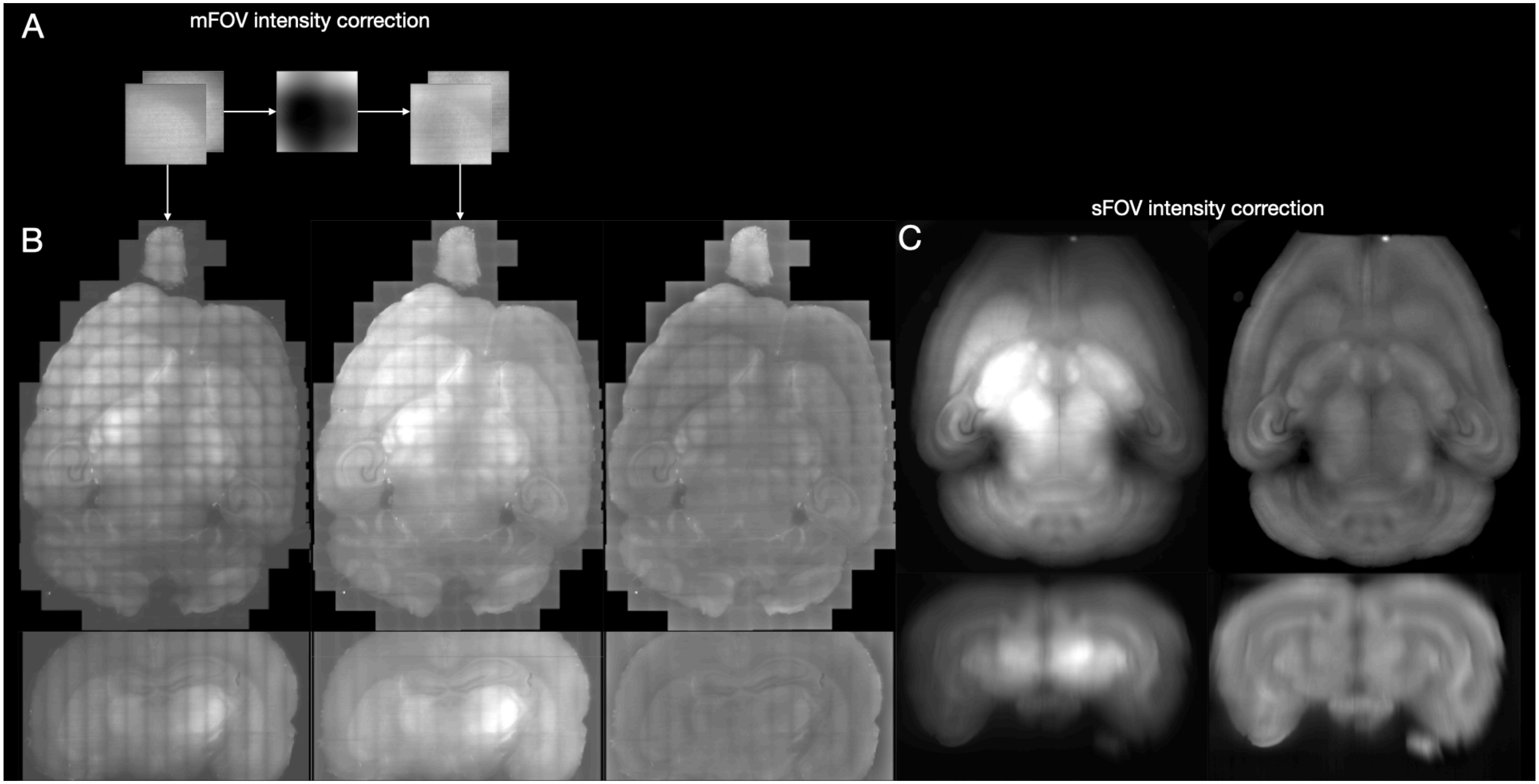


**Supplemental Figure 2**. **Intensity correction on mFOV and sFOV data.** *A:* Intensity correction per FOV. The left images show raw data from a mFOV dataset. The middle image shows the computed local multiplicative intensity correction that is applied across every FOV from the raw mFOV data. The right images show the FOVs after applying intensity correction. *B:* Effect of intensity correction on stitched mFOV data. The left column shows a single axial (top) and coronal (bottom) slice from the complete stitched volume from the raw data in A. The middle column shows the same axial (top) and coronal (bottom) slices after per-FOV correction. The right column shows the same slices after global intensity correction. *C:* Effect of intensity correction on sFOV data. Axial (top) and coronal (bottom)

slices of the raw sFOV data before (left) and after (right) global intensity correction. FOV, field-of-view; mFOV, multi-field-of-view; sFOV, single-field-of-view.

**Supplemental Figure 3**

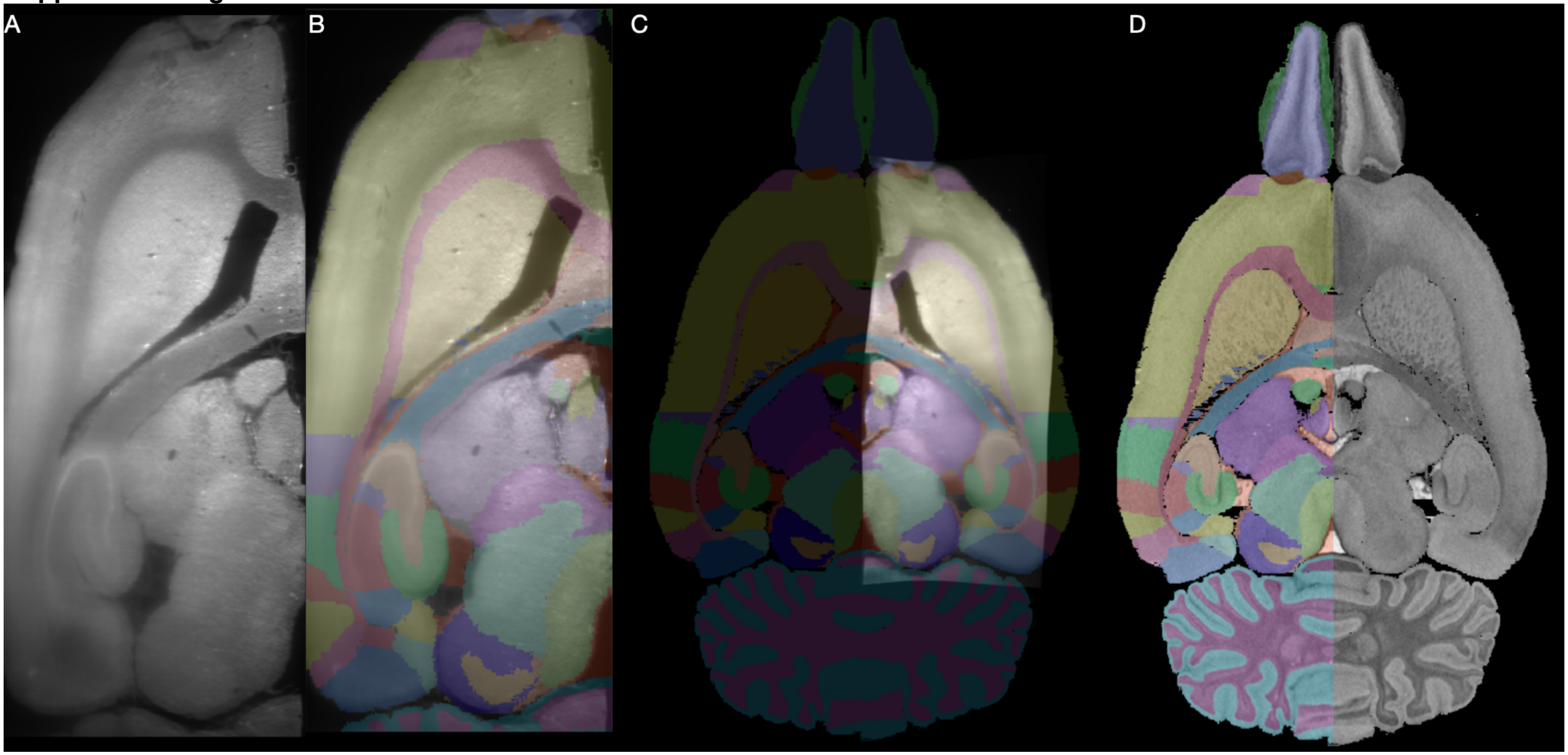


**Supplemental Figure 3: iDISCO rat hemisphere registered to Waxholm atlas.** *A:* Input data. iDISCO-cleared light-sheet microscopy-imaged rat brain region of interest. *B:* Waxholm atlas parcellations overlaid on input data. *C:* Input data transformed to Waxholm atlas. *D:* Waxholm atlas raw data with parcellations overlaid. iDISCO, immunolabeling-enabled three-dimensional imaging of solvent-cleared organs.

**Supplemental Figure 4**

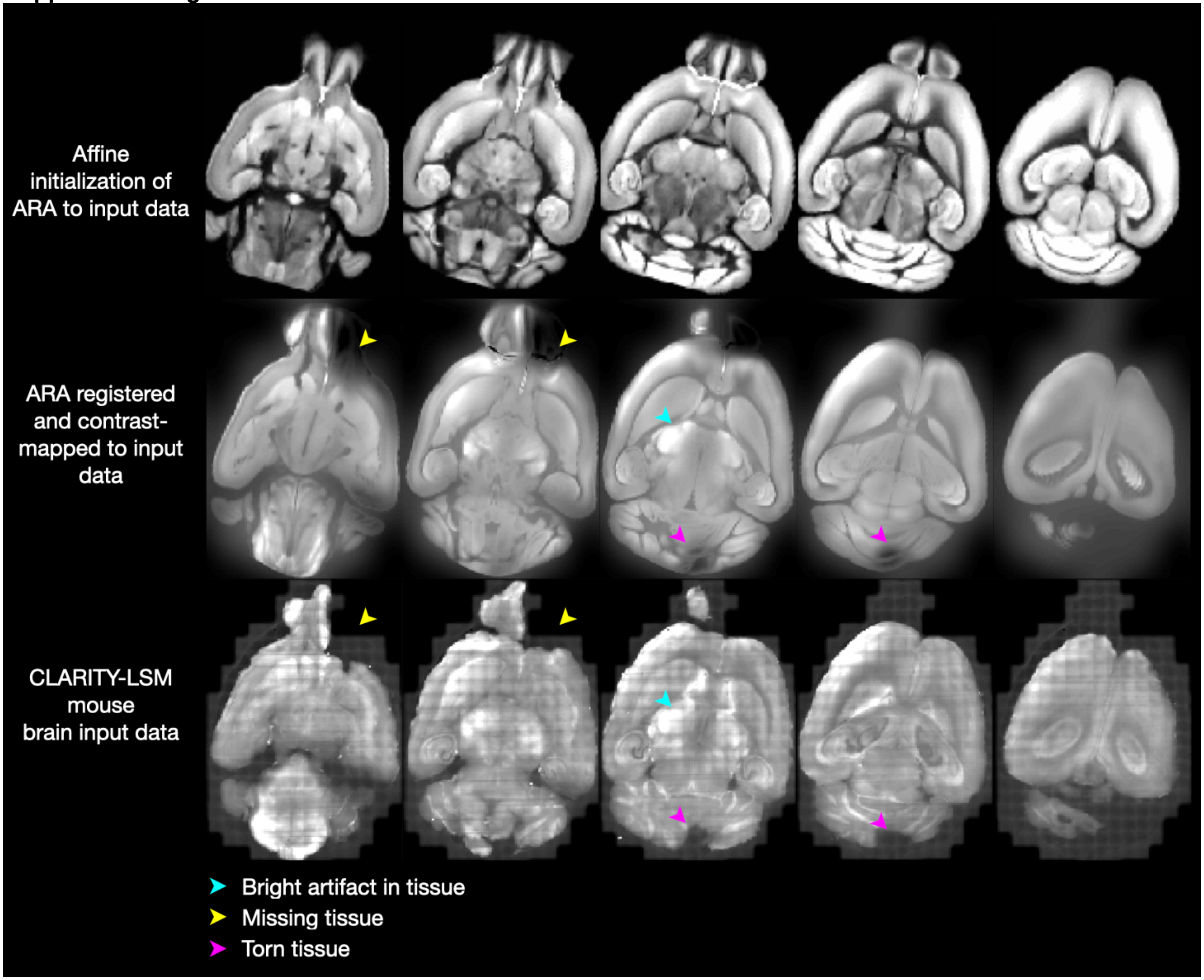

**Supplemental Figure 4: Registration and contrast-mapping tissue with artifacts.** Top row shows an affine initialization aligning the ARA CCFv3 raw data to our input data generated from LSM-imaged CLARITY cleared whole mouse brain in the bottom row. The middle row shows the ARA transformed to match the input data with a spatially varying, cubic polynomial contrast transform applied. Arrowheads indicate regions of the input tissue that contained artifacts, missing data, or torn tissue that were mapped onto the atlas using our spatially varying, cubic contrast transform, despite these significant artifacts. The bottom row contains our input data with arrowheads corresponding to the middle row. ARA CCFv3, Allen Reference Atlas Common Coordinate Frame version 3. LSM, Light-Sheet Microscopy; CLARITY, Clear Lipid-exchanged Anatomically Rigid Imaging/immunostaining-compatible Tissue Hydrogel.

**Supplementary Figure 5**

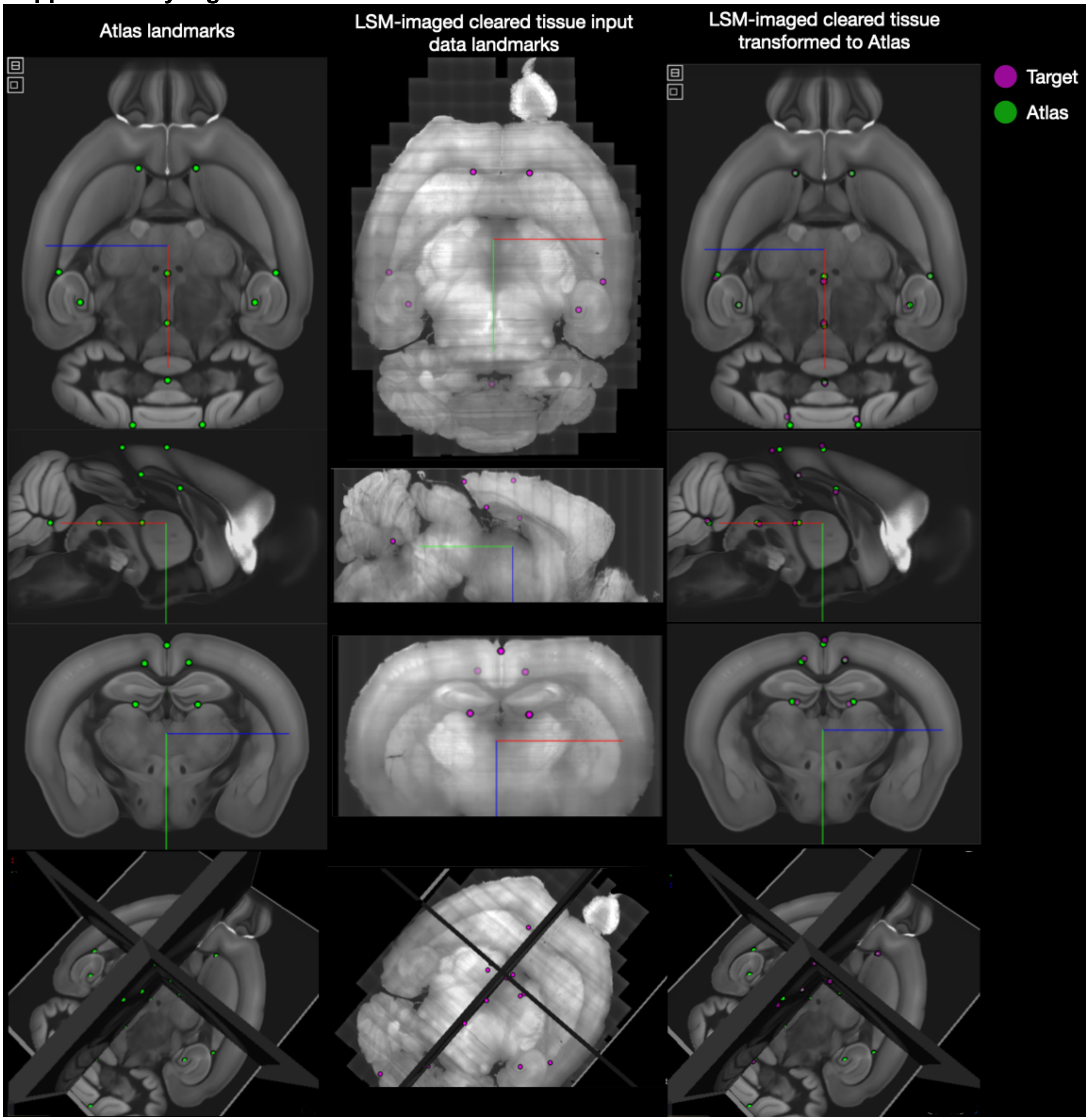


**Supplemental Figure 5: Landmarks placed for registration accuracy assessment.** *Left Column:* ARA landmarks. Locations of landmarks for assessment of accuracy shown in axial (1st row), sagittal (2nd row), and coronal (3rd row) views; landmarks are also shown in 3D space (4th row). *Middle Column:* LSM-imaged CLARTY input data landmarks. Locations of landmarks for accuracy shown in same views. *Right Column:* Input data landmarks transformed to ARA. Locations of CLARITY landmarks transformed to ARA shown in same views. CLARITY points are shown in magenta, ARA points are shown in green, and the overlap is grayscale. ARA, Allen Reference Atlas; CLARITY, Clear Lipid-exchanged Anatomically Rigid Imaging/immunostaining-compatible Tissue Hydrogel; LSM, Light-Sheet Microscopy.

**Supplemental Video 1: Registration of the Allen Reference Atlas to cleared tissue sample.** The progression of our registration over 5,000 iterations is shown. Our input LSM-imaged CLARITY mouse brain is shown in magenta and the ARA is shown in green. The overlap between the input data and the atlas shows is grayscale. LSM, Light-Sheet Microscopy; CLARITY, Clear Lipid-exchanged Anatomically Rigid Imaging/immunostaining-compatible Tissue Hydrogel; ARA, Allen Reference Atlas.

**Supplemental Table 1: CloudReg error on manually placed landmarks**

| Landmark names | Sample 1 (µm) | Sample 2 (µm) | Sample 3 (µm) | Category |
|---|---|---|---|---|
| **1** | n/a | 160 | 184 | cortex |
| **2** | 172 | 142 | 458 | cortex |
| **3** | 201 | 262 | 336 | cortex |
| **4** | 236 | 279 | 280 | cortex |
| **5** | 85 | 252 | 293 | cortex |
| **6** | 183 | 320 | 312 | cortex |
| **7** | 315 | 200 | 252 | cortex |
| **8** | 358 | 244 | 146 | cortex |
| **9** | 210 | 290 | 312 | cortex |
| **10** | 207 | 153 | 304 | cortex |
| **11** | 257 | 282 | 170 | cortex |
| **12** | 267 | 321 | 237 | cortex |
| **13** | 178 | 37 | 131 | cortex |
| **14** | 192 | 135 | 165 | cortex |
| **15** | 106 | 176 | 71 | midbrain |
| **16** | 117 | 111 | 299 | midbrain |
| **17** | 223 | 106 | n/a | cerebellum |
| **18** | n/a | 216 | n/a | cerebellum |
| **19** | 367 | 384 | n/a | cerebellum |
| **MEAN** | 227 | 242 | 274 | |
| **STDEV** | 74 | 64 | 85 | |
| **MEAN (cortex)** | 220 | 220 | 256 | 232 |
| **STDEV** | 69 | 84 | 91 | 81 |
| **MEAN (cerebellum)** | 295 | 235 | N/A | 265 |
| **STDEV** | 102 | 140 | N/A | 121 |
| **MEAN (midbrain)** | 111 | 144 | 185 | 147 |
| **STDEV** | 8 | 46 | 161 | 71 |

**Supplemental Table 1:** Euclidean distance between corresponding pairs of manually placed landmarks in the LSM-imaged CLARITY and SHIELD cleared input data and ARA is given. Mean and standard deviation (STDEV) of the landmark error is given for each sample; the mean and standard deviation of landmark error by brain region is also given for each sample. LSM, Light-Sheet Microscopy; CLARITY, Clear Lipid-exchanged Anatomically Rigid Imaging/immunostaining-compatible Tissue Hydrogel; SHIELD, Stabilization to Harsh conditions via Intramolecular Epoxide Linkages to prevent Degradation; ARA, Allen Reference Atlas.
